# Hepatoprotective Activity of Melittin on Isoniazid and Rifampicin Induced Liver Damage in Male Albino Rats

**DOI:** 10.1101/2020.09.08.287094

**Authors:** Khalid M. Naji, Bushra Y. Al-Khatib, Nora Saif Al-Haj, Myrene R. D’souza

**Affiliations:** Department of Chemistry, Faculty of Science, Sana’a University, Sana’a, Yemen; Department of Chemical Ecology/Biological Chemistry, University of Konstanz, Germany; Department of Biology, Faculty of Science, Sana’a University, Sana’a, Yemen; Department of Biochemistry, Mount Carmel College, Bengaluru, India

**Keywords:** Melittin, Bee venom, Hepatotoxicity, Tuberculosis, Rifampicin, Isoniazid

## Abstract

**Objective:** The present study aimed to investigate the ameliorative effect of melittin, a major polypeptide in the venom of honeybee (*Apis mellifera*) on isoniazid (INH) and rifampicin (RIF) induced hepatotoxicity in male albino rats. *Method*: The rats (140-200g) were divided into five groups (n=6): normal control (NC); toxic (T) group treated with INH+RIF (100 mg/kg, p.o.); melittin-treated (Mel15, Mel30) group (15 or 30 µg/kg s.c); and normal recovery (NR) group. Blood and liver samples were collected for biochemical, hematological and histopathological studies respectively.

**Results:** The administration of melittin was found to prevent the antitubercular drug-induced alterations in the diagnostic markers; reduced glutathione (GSH), direct bilirubin (DB), total bilirubin (TB), aspartate aminotransferase (AST), alanine aminotransferase (ALT), lactate dehydrogenase (LDH), alkaline phosphatase (ALP), and total serum protein (TSP). Besides, hematological alterations were significantly high (P<0.0001) in Mel-treated groups when compared to the toxic control. The NR group exhibited lower levels of DB, TB, ALP, LDH and TSP. In addition, treatment with melittin offered protection in the NR group with respect to MDA levels.

**Conclusion:** Evidence from this study indicate that melittin is beneficial for the prevention of acute hepatic failure in antitubercular drug-induced hepatoxicity.

## Introduction

Despite tremendous advances in modern medicine, the prevention and treatment of liver diseases such as chronic hepatitis, cirrhosis, fibrosis, and hepatocellular carcinoma remain serious health complications in the world today [1]. The unique metabolic function and relationship of the liver to the gastrointestinal tract makes it an important target of toxicity to xenobiotics, hepatotoxins, environmental pollutants and absorbed drugs [2]. Certain drugs may be hepatotoxic as primary compounds even when administered within the therapeutic ranges; or may form reactive metabolites or trigger immune-mediated responses that effect hepatocytes, biliary epithelial cells and/or liver vasculature [3,4].

Tuberculosis is a highly communicable disease affecting over one-third of the world’s population and killing over 2 million people per year [5]. A meta-analytical study on the co-administration of isoniazid (INH) and rifampicin (RIF), the first line drugs used for tuberculosis therapy is associated with 2-6% hepatotoxicity [6], acute liver injury and high mortality rate [7]. These drugs mediate the generation of highly reactive oxygen species (ROS), which act as a trigger of lipid peroxidation and a source for destruction to the plasma membrane [8]. RIF actively induces CYP2E1, a member of the cytochrome P450 family responsible for the breakdown of environmental chemicals and carcinogens. It also increases INH induced toxicity by stimulating the formation of the toxic metabolite hydrazine when it proceeds along the amidase pathway. Hydrazine then reacts with sulfhydryl group of glutathione (GSH), depletion its levels within the hepatocytes and causing cell death [2]. Combination chemotherapy with INH-RIH was found to stimulate the conversion of INH to isonicotinic acid, another hepatotoxic product. The plasma half-life of acetyl hydrazine is further shortened by RIF as it is quickly converted to its active metabolites; thereby increasing the occurrence of liver necrosis [9]. Thus, administration of RIF is correlated to an increase in the incidence of anti-TB drug - induced hepatotoxicity [10].

Many traditional remedies employing herbal drugs continue to make a major contribution to health in terms of prevention and treatment of many diseases [11]. Venomous insects have long been used in scientific research and represent the basis of many traditional drugs [12]. Bee venom, a natural toxin produced by the honeybee (*Apis mellifera*) is known to relieve pain and treat inflammatory diseases, such as rheumatoid arthritis, in both humans and experimental animals [13,14]. Bee venom comprises of a variety of peptides, including melittin, adolpin, apamin, phospholipase A2 and mast cell degranulating peptide [15]. Melittin (MEL) is the principal toxin of bee venom comprising approximately 50% of its dry matter. MEL is a small linear basic peptide with the chemical formula C_131_H_228_N_38_O_32_, twenty-six amino acid long and weighs 2847.5 Da [16]. It suppresses inflammation by inhibiting phospholipase (PLA) activity [17]. The protective effect of MEL on various inflammatory parameters has been reported by several investigators on acute liver inflammation [18], osteoarthritic chondrocytes [19], rheumatoid arthritis [20], and acute pancreatitis [21]. MEL was found to attenuate inflammation and fibrosis by inhibiting the NF-κB signalling pathway in liver fibrosis induced by thioacetamide. It decreases the rate of lethality, inhibits hepatocyte apoptosis and attenuates hepatic inflammatory responses. Recent reports in various disease models have proved the anti-inflammatory effects of MEL [22]. In the current study, we aimed to assess the effects of melittin on anti-tuberculosis drugs-induced hepatotoxicity via biochemical analysis, haematological and histological examination.

## Materials and Methods

### Chemicals

Rifampicin was procured from Himedia Laboratories, India. Melittin, GSH, DTNB, TBA, methanol and ethanol were purchased from Sigma-Aldrich Chemical Company (St. Louis, MO, USA). Isoniazid was procured from Wako Pure chemical industries Ltd.

### Experimental animals

The study was carried out according to the guidelines prepared by the National Academy of Sciences and published by the National Institute of Health (*Guide for the Care and Use of Laboratory Animals, 1996*) [23]. The Committee of Experimental Animals Care and Use, Sana’a University approved the protocol followed in this study.

For this experiment, thirty adult male rats *Rattus rattus*, weighing 140-200 g were collected from the animal house, Biology Department, Faculty of Science, Sana’a University, Sana’a. The animals were housed in the research laboratory, Department of Pharmaceutics, University of Science and Technology, Sana’a in separate standard plastic cages (0.90 × 0.60 × 0.40 m) and maintained under room temperature with alternating 12 hours light/dark cycle. The animals were provided with food pellets and water *ad libitum*. Collection trays were placed below the cages. The amount of food and water taken by the rats were recorded every day, while the body weight was recorded every week.

### Experiment design

After the adaptation period, the rats were randomly divided into five groups of six animals:

i. ***Normal control (NC) group*:** Rats received normal saline (NaCl, 0.9%) daily, throughout the experimental period, without any treatment.
ii. ***Toxic (T) group*:** Rats were treated with INH (100 mg/kg body weight, p.o.), and co-administered with RIF (100 mg/kg body weight, p.o.) for 21 days.
iii. ***Mel15 group*:** Rats were treated with INH (100 mg/kg body weight, p.o.), and co-administered with RIF (100 mg/kg body weight, p.o.) for 21 days + MEL (15 μg/kg MEL, subcutaneously) for 15 days.
iv. ***Mel30 group*:** Rats were treated with INH (100 mg/kg body weight, p.o.), and co-administered with RIF (100 mg/kg body weight, p.o.) for 21 days + MEL (30 μg/kg MEL, subcutaneously) for 15 days.
v. ***Normal recovery (NR) group***: Rats were treated with INH (100 mg/kg body weight, p.o.), and co-administered with RIF (100 mg/kg body weight, p.o.) for 21 days. The animals were then allowed to recover for 15 days without any treatment.

### Preparation of the samples for biochemical studies

After overnight fasting, two sets of blood samples were collected via the retro-orbital venous plexus using microhematocrit capillary tubes: first, into an EDTA containing tube, to obtain plasma for the hematological studies; and second, into a tube with no additives to obtain serum for biochemical analysis. All rats were then sacrificed under ether anesthesia, the liver was dissected out, weighed and washed immediately with ice-cold saline to remove as much blood as possible. Each liver sample was divided into two parts - one part was fixed in 10% neutral-buffered formalin for histology analysis, while the second part was used for preparing crude extracts. 10% extracts were prepared by homogenizing the tissue in ice-cold 0.1 M phosphate buffer (pH 7.4) with Teflon homogenizer at 3000 rpm for 10 min. The homogenate was centrifuged at 15,000 rpm for 30 min followed by lyophilization of the supernatant. The supernatant served as the source for reduced glutathione (GSH), lipid peroxidation (LPO) and total protein (TP). The preparation of tissues samples was carried out at the Tropical Disease Research Center–University of Science and Technology Hospital.

### Measurement of serum ALT, AST, ALP, TB and TSP

Serum biochemical parameters such as direct bilirubin (DB), total bilirubin (TB), aspartate aminotransferase (AST), alanine aminotransferase (ALT), lactate dehydrogenase (LDH), alkaline phosphatase (ALP), and total serum protein (TSP) were determined using kits supplied by Roche Diagnostics and the Roche/Hitachi Analyzer machine at Al-Aulaqi Specialized Medical Laboratory, Sana’a.

### Biochemical analysis of tissues

The following tests were performed for biochemical analysis of tissues.

#### Lipid Peroxidation (LPO)

Malondialdehyde (MDA) levels were estimated by its reaction with thiobarbituric acid [24] to form a complex called thiobarbituric acid reactive substance (TBARS) that absorbs at 535 nm. The values were expressed as nmol/mg tissue.

#### Reduced glutathione (GSH)

Reduced glutathione reacts with 5,5′-dithiobis(2-nitrobenzoic acid) to produce a yellow colored product with absorption maxima at 412 nm [25]. A calibration curve was plotted with standard GSH (2-10 nmole) and values obtained was expressed as nmole GSH/mg tissue.

#### Total protein

Total protein (TP) contents of liver, heart and aorta homogenates was determined at 660 nm [26].

### Hematological assays

The red blood cells (RBCs), total white blood cells (T-WBCs) count, differential leukocytes count, platelets, hemoglobin (Hb), hematocrit (Hct), mean corpuscular hemoglobin (MCH), mean corpuscular hemoglobin concentration (MCHC), mean corpuscular volume (MCV), and red cell distribution width (RDW-CV) were measured using automated Hematology System, Sysmex XT-2000i at Al-Aulaqi Specialized Medical Laboratory, Sana’a.

### Histological analysis

#### Preparation of tissue samples

After the experimental period, all the rats were dissected and the liver tissue was removed and washed in ice normal saline solution, fixed in 10% formalin, dehydrated in graded alcohol (50% -100%) and embedded in paraffin. The tissue was cut (3 µm) and stained with hematoxylin and eosin [27]. Sections were examined under a light microscope (Olympus microscope CX-21) and photographed by attached Olympus Camera DP21 (U-TV0.5XC-3).

#### Histopathological quantitative analysis

Quantitative analysis of the histopathological changes in the liver was averaged (n=6 per group) using an ocular micrometer calibrated with a stage micrometer. The frequency of histopathological changes was based on an average obtained from an observation of 10-microscopic fields [28] with an area 625μm^2^ at 40x or 400x. The histology work was performed in the Histology Lab, at the University of Science and Technology.

### Statistical analysis

All data presented were expressed as mean ± SD. One-way analysis of variance (ANOVA) was used for analyzing statistical significances between the groups. This was followed by Tukey Multiple Comparisons with the help of Prism 6 software (Graph Pad, San Diego, CA, USA). Values with P < 0.05 was considered significant.

## Results

### Body and Liver weight

At the end of the experiment, there were no significant changes in the bodyweight of the animals (Table 1). The T group showed a non-significant decrease compared to the NC group, whereas there was a significant increase in Mel 30 group compared to the T group (P<0.1).

**Table 1.** Body weight and liver weight of male rats treated with melittin post isoniazid and rifampicin-induced hepatotoxicity

|  | NC | T | Mel15 | Mel30 | NR |
| --- | --- | --- | --- | --- | --- |
| **Body weight (g)** | 184.2± 18.3 | 174.7±24.3 | 209±20.5 | 220.5±18.1 <sup>b*</sup> | 191.8±41.6 |
| **Liver weight (g)** | 9.788±1.05 | 7.57±1.26 | 10.26±0.99 | 11.28±2.38 <sup>b**</sup> | 9.08±2.23 |

Results are expressed as mean ± SD (n=6). NC: control, T: Toxic, Mel15: treated by 15µg Melittin,
Mel30: treated by 30µg Melittin, NR: Normal recovery. \* $P<0.05$ ; \*\* $P<0.01$ ; b: significant Vs T group

### Biochemical markers

Rats in the T group treated with anti-tubercular drugs exhibited a significant increase in the levels of DB, TB, LDH (P< 0.0001), ALT, ALP (P<0.001), AST, TSP (P<0.01) compared to NC group (Table 2). Conversely, the Mel15 group displayed a significant decline in the levels of DB, TB, ALP (P < 0.0001), ALT (P<0.01), AST (P<0.05) and LDH (P<0.001) compared to the T group. Similarly, there was a significant decrease in DB, TB, ALP (P<0.0001), ALT, AST (P<0.05) and LDH (P<0.001) levels in the Mel30 group when compared to the PC group. There was no significant change in TSP when compared to group T. On the other hand, there was a significant decrease in serum DB, TB (P<0.0001), ALP, TSP (P<0.001) and LDH (P<0.05) in the NR group when compared with the T group (Table 2).

**Table 2.** Biochemical parameters of male rats treated with melittin post isoniazid and rifampicin-induced hepatotoxicity

|  | NC | T | Mel15 | Mel30 | NR |
| --- | --- | --- | --- | --- | --- |
| **DB (μmol/l)** | 1.0±0.451 | 7.05±1.84<sup>a****</sup> | 1.0±0.38<sup>b****</sup> | 1.0±0.27<sup>b****</sup> | 1.0±0.31<sup>b****</sup> |
| **TB (μmol/l)** | 1.6±0.94 | 10.45±2.13<sup>a****</sup> | 2±1.05<sup>b****</sup> | 2.82±1.49<sup>b****</sup> | 2.87±0.89<sup>b****</sup> |
| **ALT (U/L)** | 26.83±11.44 | 73.5±16.99<sup>a***</sup> | 38.5±14.27<sup>b**</sup> | 44.33±15.93<sup>b*</sup> | 50.33±17.44 |
| **AST (U/L)** | 92.17±31.64 | 288.3±147.7<sup>a**</sup> | 137.7±15.81<sup>b*</sup> | 147±32.83<sup>b*</sup> | 183.2±79.05 |
| **ALP (U/L)** | 187.2±25.86 | 286.5±53.07<sup>a***</sup> | 164.2±19.59<sup>b****</sup> | 166.5±18.41<sup>b****</sup> | 189±27.41<sup>b***</sup> |
| **LDH (U/L)** | 762±63.68 | 3046±768.1<sup>a****</sup> | 1577±338.4<sup>b***</sup> | 1640±693.3<sup>b***</sup> | 2155±261.2<sup>b*</sup> |
| **TSP (g/L)** | 66±8.37 | 75.5±0.84<sup>a**</sup> | 76±1.67 | 70±2.76 | 63.5±2.51<sup>b***</sup> |

Results are expressed as mean ± SD (n=6). NC: control, T: Toxic, Mel15: Melittin15μg, Mel30: Melittin30μg, NR: Normal recovery. \*P<0.05; \*\*P<0.01; \*\*\*P<0.001; \*\*\*\*P<0.0001.
a: significant Vs NC group, b: significant Vs T group

### Antioxidant markers and total protein

#### Lipid peroxidation (LPO)

The extent of lipid peroxidation analysed by MDA levels, indicated a significant elevation in the T group when compared with the NC group. A significant increase in MDA was demonstrated in NR (P<0.0001) when compared with T group. On the other hand, there was an insignificant reduction in liver MDA in Mel15 and Mel30 groups compared with the T group as shown in Figure 1A.

**Figure 1:**
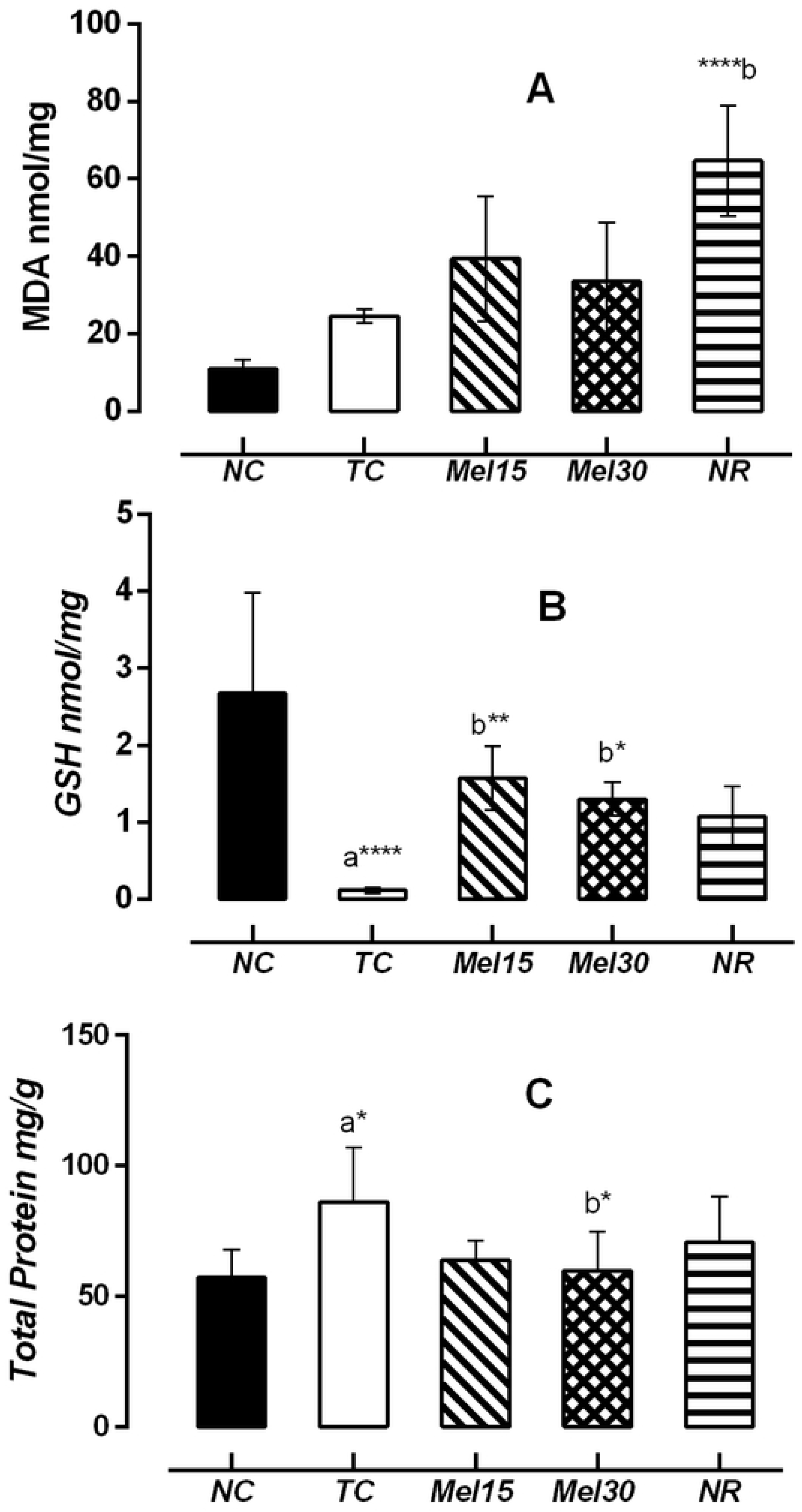
Effect of Melittin in the levels of MDA, GSH and TP in post isoniazid and rifampicin-induced hepatotoxicity. Results are expressed as mean ± SD for at least six rats per group. NC: control; T: Toxic; Mel15: Melittin15µg; Mel 30: Melittin30µg; NR: Normal recovery; *\*P<0*.*05; **P< 0*.*01; ***P<0*.*001; ****P<0*.*0001*. a: significant vs NC group, b: vs T group

#### Reduced glutathione (GSH)

The GSH content in liver of T group declined in comparison to the NC group (P<0.0001). Furthermore, a significant increase in GSH level was observed in the Mel15 (P<0.01) and Mel30 (P<0.05) groups compared with the T group after 15 days of treatment. No significant difference in GSH level was observed in the NR group compared to the T group (Figure 1B).

#### Total protein (TP)

Total protein levels in liver showed a significant increase in T group (P<0.05) when compared to NC group, while there was a significant decrease in Mel30 group (P<0.05) when compared with T group. No significant changes between the rats treated with Mel15 and NR group was seen (Figure 1C).

#### Hematological parameters

Compared to NC group, the administration of INH +RIF caused a significant decrease in NEU, RBCS, Hb, PLT (P <0.01), MCV and RDW-CV (P< 0.0001), Hct, MCH (P< 0.001) while BAS, MCHC was increased significantly (P< 0.0001) (Table 3). WBCs (P< 0.0001), PLT (P< 0.001), Hb, Hct, MCV (P < 0.01), EOS (P < 0.05) rose significantly. After administration of MEL for 15 days, a significant decline in BAS and MCHC (P< 0.0001) when compared to T group was reported. A significant increase was found in Mel30 group for WBCs (P< 0.0001), NEU, Hb, PLT and RDW-CV (P < 0.01), Hct and MCV (P< 0.001), EOS and MCH (P< 0.05), while BAS and MCHC were significant decreased (P< 0.05) in comparison with T group. On the other hand, there was a significant increase in WBCs and Hb (P < 0.05), NEU (P < 0.001), EOS, Hct, MCH (P< 0.01) and MCV (P< 0.0001). There a significant decrease in BAS, MCHC (P< 0.0001) in the NR group compared to the T group. Moreover, LYM and MON in groups NC, T, Mel15, Mel30 and NR had no statistical changes.

**Table 3.**
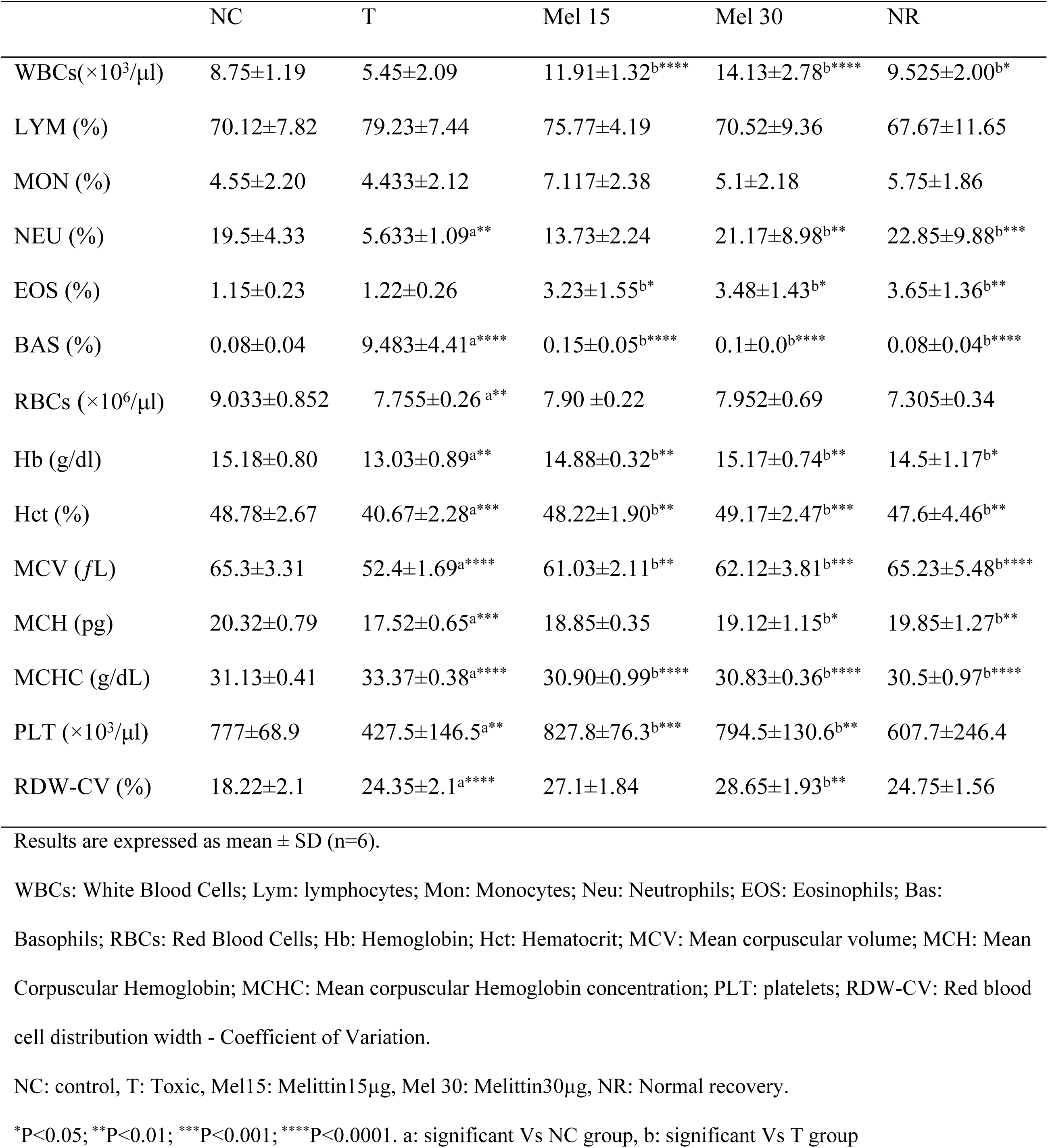
Complete blood count in male rats treated with melittin post isoniazid and rifampicin-induced hepatotoxicity

#### Histopathology

Histopathological examination provided important evidence supporting the results obtained from biochemical tests. In this study, quantitative analysis for histological lesions was used. Liver sections of control group exhibited a central vein and normal hepatocytes with a well-preserved cytoplasm and a prominent nucleus and nucleolus as shown in Figure 2a. Liver sections of hepatotoxic group revealed severe histopathological lesions as shown in Table 4 and Figure 2 (b, c, d & e). In the quantitative assessment, all scores of hepatic histopathological lesions in the toxic group were shown to be significantly higher than the NC group. Lesions demonstrating hemorrhage, congestion, ballooned hepatocytes, cellular inflammation, necrosis, fibrosis, amyloids (P< 0.0001) and vasodilation (P<0.001) indicated that INH+RIF caused severe hepatic damage. On the other hand, Mel15 group demonstrated significantly lower scores of hemorrhage and congestion (P< 0.001), ballooned hepatocytes and inflammation (P< 0.01), while the greatest decrease was detected for necrosis, fibrosis and amyloids (P< 0.0001) when compared to T group as shown in Table 4 and Figure 2f.

**Table 4.** In situ evaluation of rat’s liver treated with melittin post isoniazid and rifampicin-induced hepatotoxicity

|  | NC | T | Mel15 | Mel30 | NR |
| --- | --- | --- | --- | --- | --- |
| **Amyloids** | - | ++++ | - | - | + |
|  | 1.33±0.52 | 17.5±6.47<sup>a****</sup> | 2.67±1.21<sup>b****</sup> | 1.5±0.55<sup>b****</sup> | 3.83±1.72<sup>b****</sup> |
| **Congestion** | + | ++++ | + | + | +++ |
|  | 1.33±0.52 | 6.67±1.36<sup>a****</sup> | 2.4±1<sup>b***</sup> | 2.17±0.98<sup>b***</sup> | 5.17±2.56 |
| **Ballooned hepatocytes** | - | ++++ | + | ++ | ++ |
|  | 4±2 | 86±39.43<sup>a****</sup> | 30±11.65<sup>b**</sup> | 49.4±17.81 | 54.3±19.79<sup>a**</sup> |
| **Inflammation** | - | ++++ | ++ | ++ | ++++ |
|  | 0.83±0.41 | 4.5±0.84<sup>a****</sup> | 2.5±1.0<sup>b**</sup> | 2.5±1.2<sup>b**</sup> | 4.4±1.0 |
| **Necrosis** | + | ++++ | + | + | +++ |
|  | 13.17±4.8 | 64.83±17.2<sup>a****</sup> | 23.17±10.1<sup>b****</sup> | 22.33±4.1<sup>b****</sup> | 43.5±14.3<sup>b*</sup> |
| **Vasodilation** | - | ++++ | ++ | +++ | ++++ |
|  | 1.8±0.41 | 16.3±4<sup>a***</sup> | 11.2±4.7 | 12.5±4.76 | 20.3±7.2 |
| **Fibrosis** | - | +++ | - | + | ++++ |
|  | 0.83±0.41 | 6.83±2.0<sup>a****</sup> | 1.3±0.5<sup>b****</sup> | 1.8±0.4<sup>b***</sup> | 7.3±3.2 |
| **Hemorrhage** | - | +++ | + | ++ | ++++ |
|  | 1.5±0.55 | 5.5±0.55<sup>a****</sup> | 2.13±0.95<sup>b***</sup> | 4.5±2.1 | 9.167±1.17<sup>b***</sup> |
| **Steatosis** | - | + | - | - | ++++ |
|  | 1.3±0.5 | 16.5±5.1 | 3±1.1 | 6.67±1.6 | 48.5±19.9<sup>b****</sup> |

Results are expressed as mean ± SD (n=6).
NC: control, T: Toxic, Mel15: Melittin15µg, Mel 30: Melittin30µg, NR: Normal recovery
Damages graded as follows: -, absent; +, trace (1–25%); ++, mild (25–50%); +++, moderate (50–75%); +++++, severe (75–100%). \*P<0.05; \*\*P<0.01; \*\*\*P<0.001; \*\*\*\*P<0.0001.
a: significant Vs NC group, b: significant Vs T group

**Figure 2.**
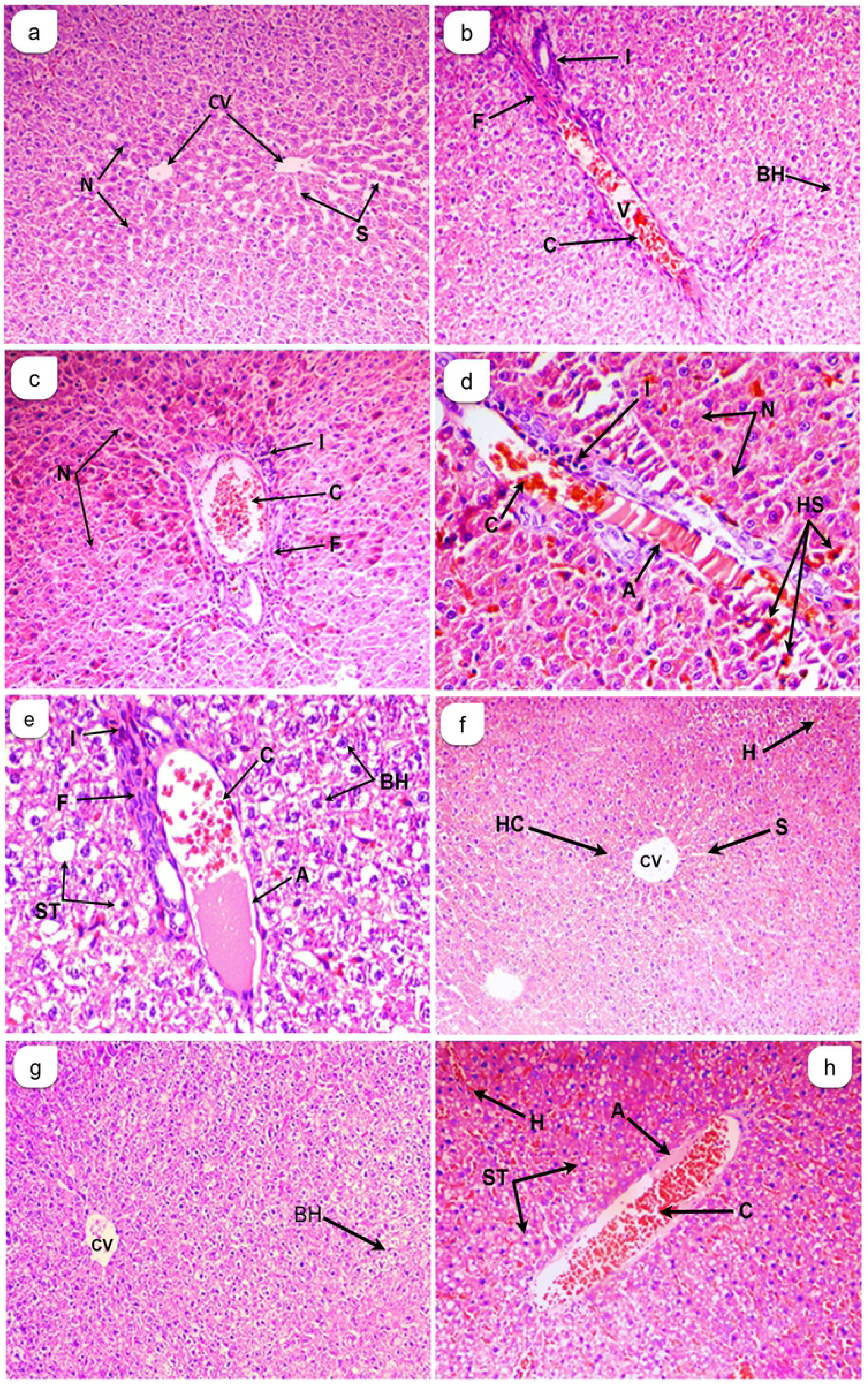
Cross sections in liver of rats showed the effect of Melittin on liver tissues in post isoniazid and rifampicin-induced hepatotoxicity. (a): Normal control (NC); (b): toxic group (T); (c): treated group 15µg Melittin; (d): treated group 30µg Melittin (Mel 30); (e): Normal recovery group (NR). (a): Normal group (N), 200x; (b & c): Toxic group (T), 200x; (d & e): Toxic group(T) 400x; (f): Mel 15 mg, 200x; (g): Mel 30mg, 200x; (h): Normal recovery group(NR) 200x. *Histological abbreviations*: Amyloids (A), Congested blood vessel (C), Ballooned hepatocytes (BH), Inflammation (I), Necrosis (N), Vasodilation (V), Fibrosis (F), Hemorrhage (H), Steatosis (ST), Hepatocytes (HC), Central vein (CV), Sinusoid (S) and Hemosiderin-ladenmacrophages (HS)

Furthermore, Mel30 group had significantly decreased scores for congestion and fibrosis (P< 0.001), and inflammation (P< 0.01). The greatest decrease was however detected with necrosis and amyloids (P<0.0001) as compared to T group (Table 4 and Figure 2g). In addition, the NR group had significantly decreased scores of hemorrhages (P< 0.001), and steatosis (P< 0.05), while showing significant reduction in necrosis (P< 0.05), and amyloids (P< 0.0001) when compared with T group (Table 4 and Figure 2h).

## Discussion

The present investigation examines the protective effects of the peptide, melittin a major component of bee venom against hepatotoxicity induced by the co-administration of Isoniazid (INH) + Rifampicin (RIF) in adult male albino rats for 21 days. The weight of the body and liver of rats treated with INH+RIF was insignificantly lower than the NC group. This is in agreement with previously documented hepatotoxicity studies that attribute the decrease in food intake to anorexia and reduced food palatability manifested on account of drug-induced toxicity [29,30]. INH+RIF induces oxidative stress by generating free radicals and altering the antioxidant status resulting in metabolic disorder and weight loss [29]. A dose-weight dependent increase in both body and liver weight was seen on treatment with MEL (15, 30 µg/kg) when compared the T group; with Mel30 being more effective than Mel15 and NR groups. Thus, it could be implicated that treatment with MEL improves the food intake, food palatability, metabolic processes, and consequently the body weight and liver/body weight ratio of INH+RIF treated rats.

Co-administration of INH+RIF for 21 days caused a significant increase in the serum levels of DB, TB, ALT, AST, ALP, LDH and TSP. The findings of this study are in agreement with numerous reports that document a rise in the activity of these serum enzymes [31–33]. However, the exact mechanism of hepatotoxicity by INH+RIF is still not clear, although some researchers have suggested that hepatotoxicity is mediated through release of reactive/toxic metabolites that bind covalently to the macromolecules of hepatocytes [34].

Bilirubin, a product of RBC breakdown is a good indicator of liver and bile duct function [35]. However, hyperbilirubinemia seen in acute hepatitis is directly proportional to the degree of histological injury of the liver and duration of the disease [36]. The combination of INH+RIF resulted in a higher rate of inhibition of biliary secretion, an increase in lipid peroxidation in the hepatocyte, and cytochrome P450 involved in the synergistic effect of RIF and INH [37].

Increase in activity of the enzymes ALT and AST is indicative of massive hepatocellular disease and is linked to acute hepatic necrosis [38]. Chronic liver disease is diagnosed by a large increase of mitochondrial AST in the serum following extensive tissue necrosis [39]. MEL treatment reduced the elevation of AST and ALT activities in GalN/LPS-induced liver injury [40]. Similarly, pre-treatment of EAT-bearing mice with MEL (0.5, 5 or 50 µg/100 g of body weight, every other day for 4 weeks) caused a progressive decrease in serum levels of the liver enzymes when compared with those of the EAT-bearing control mice [41]; with MEL groups showing greater improvement than the NR group.

The increase in ALP activity (Table 2) is related to improper bile flow through the gall bladder and bile duct into the intestine [42]. Leaky tight junctions found between the canaliculi wall cells could cause leakage from the bile canaliculi into hepatic sinusoids [43]. Furthermore, overproduction and release of ALP into the blood due to hepatobiliary injury and cholestasis may also contribute to an increase in serum ALP activity [37]. On the other hand, the high serum levels of LDH, an important index of liver cell damage may arise due to hepatocellular necrosis. Similar results were reported by [44]. In was hence noted that MEL at different levels had a significant effect on liver enzyme markers and could contribute to protecting liver epithelial cells from damage through the inhibition of inflammatory cytokines and apoptosis [18,45]. It could be suggested that the administration of MEL may prevent liver damage related to the maintenance of plasma membrane integrity, thereby suppressing the leakage of enzymes through membranes. Additionally, MEL may also exhibit hepatocurative activity against the hepatotoxicity induced by combination of INH and RIF drugs.

Anti-tubercular drugs through oxidative stress causes cellular damage as well as dysfunction of the hepatic antioxidant defense system [46]. Enhanced susceptibility of the hepatocyte cell membrane to anti-tubercular drugs induce peroxidative damage which increases the activity of diagnostic marker enzymes (Table 2) in the systemic circulation. In our study, we found there was variation in T group for levels of MDA, GSH and TP. Increased levels were observed in MDA and TP, while a significant decrease in GSH when compared to NC group was reported (Figure 1). The effect of free radicals formed either by the reaction of metabolites of INH/RIF with oxygen or by the interaction of superoxide radicals with H_2_O_2_ seems to initiate peroxidative degradation of membrane lipids and endoplasmic reticulum rich in polyunsaturated fatty acids [47]. A statistically significant (P< 0.0001) increase in MDA in NR group when compared to T group (Figure 1) reflect hepatocellular damage and the depletion of antioxidant defenses or that the raising in free radical production deteriorates the prooxidant-antioxidant balance leading to oxidative stress-induced cell death [48].

The depletion of reduced GSH (Figure 1) is known to be the result of enhanced lipid peroxidation and hence an increase in glutathione consumption [49]. The depletion of hepatic GSH in toxic conditions provides useful information on the protective role of GSH against toxic foreign compounds or drugs [50]. The decline in GSH levels was also reported by other researchers [36,51] and can be ascribed to the biotransformation of INH and RIF by glutathione conjugation [52]. Besides, the levels of GSH in Mel15 and Mel30 groups increased significantly providing evidence for its anti-inflammatory role. These findings were in partial similarity with results in GalN/ LPS-induced liver injury [40].

The significant decline in hematological parameters NEU, RBCs, Hb, Hct, MCV, MCH and PLT level in T group when compared to NC group; while a significant increase in MCHC and RDW-CV are in agreement with INH and RIF induced hepatotoxicity by many researchers [53]. Anti-tubercular drugs have been recorded to cause hemolytic and aplastic anemia by the depression of the bone marrow [54,55]. INH and its metabolite in particular affect the erythroid precursor cells, the formation of autoantibodies and inhibition of DNA synthesis [56,57]. Furthermore, INH binds to pyridoxine (vitamin B6), depleting its levels and causing anemia [58]. The severity of neutropenia associated with drugs is often dose-dependent, but the occurrence of reactions is said to be idiosyncratic and agranulocytosis associated with direct toxicity is usually associated with a slower decline in neutrophil levels [59].

The result of this study revealed a decrease in Hb level (Table 3) probably due to INH-mediated inhibition of heme synthesis by depressing δ-aminolevulinic acid (ALA) synthetase, the first and rate-limiting enzyme in the heme synthesis pathway [60]. The decrease in MCV and MCH are indicative of microcytic hypochromic anemia, while the observed increase in MCHC may cause hemolysis marked to spherocytosis, which was affirmed in previous findings [61]. The significant decrease in platelets as seen in Table 3 could result in thrombocytopenia as was also reported by other researchers working on anti-tubercular drugs [62]. Thrombocytopenia occurred mostly due to RIF binding non-covalently to thrombocyte membrane glycol-proteins IIb/IIIa or Ib/IX to produce compound epitopes or induce conformational changes [63]. The mechanism by INH induced thrombocytopenia is still unclear. An increase in BAS and RDW-CV in T group when compared to the NC group (Table 3) is seen. Basophil count significantly raising is very rare, but if seen, indicates myeloproliferative disorder or other more obscure causes. On the other hand, a greater decrease in Hb in patients with higher values for RDW was reported [64]. In iron deficiency anemia (anisocytosis), the varied size distribution of RBC causes an increase in the RDW [65].

In the Mel15 and Mel30 groups, a significant increase in WBCs, NEU, EOS, Hb, Hct, MCV, MCH and PLT, while a significant decrease in BAS compared to T group was observed (Table 3). The same results were also seen in the NR group but the improvement of the studied hematological parameters in the MEL groups was higher than that in the NR group. This may suggest a positive effect of MEL against blood coagulation, triggering of erythrocytes formation and the regeneration of leucocytes and erythrocytes [66]. The increasing of the EOS in Mel15 and Mel30 groups may be due to the allergenic natural of melittin. This result was affirmed when several patients taking MEL complained from allergy symptoms [67]. Other studies reported that the MEL plays a central role in the production of nociceptive responses and cutaneous hypersensitivity after whole bee venom injection [68,69].

Our histopathological findings in liver sections such as amyloids, congestion, ballooned hepatocytes, cellular inflammation, necrosis, vasodilation, fibrosis, hemorrhage were statistically evaluated. The marked changes occurring in liver tissues support the results obtained by studies on serum markers. Amyloids deposits were found to be the highest in the T group (Table 4). These are aggregations of misfolded proteins that trigger tissue injury and impair normal cellular function due to the exertion of pressure on hepatic tissue [70]. This explain by the appearance of a high amount of TP in the tissue, resulting in the emergence of the case of amyloid in the T group. Vasodilation observed to be the highest in T group (Table 4) may have been induced by the action of several mediators notably histamine on vascular smooth muscle. Vasodilation leads to increased permeability of the microvasculature, causing the outpouring of protein-rich fluid into extravascular tissues. The vessels diameter is increased leading to slower blood flow, increased concentration of red blood cells in small vessels, and increased viscosity of the blood causing engorgement of the small vessels. This condition termed as stasis was seen as vascular congestion and localized redness of the involved tissue. Number of ballooned cells in the liver of the T group was found to be the highest among all groups (Table 4). These cells are formed by excess water accumulation in the vacuoles as a result of the failure of active membrane transport due to loss of ATP as the energy source [71].

Toxic mediators generated by phagocytes aggravates liver injury causing inflammation [72]. The transcription factor NF-*κ*B, upon translocation into the nucleus, expresses inflammatory mediators implicated in the regulation of genes related to apoptosis, inflammation, and immune response [73]. RIH triggers cytokine-induced production of pro-inflammatory mediators such as NO and IL-8, in the hepatocyte cell line [74]. Co-administration of INH+RIF induces a significant increase in serum levels of the pro-inflammatory cytokine TNF-α, a mediator of inflammatory tissue damage [75]. In our study, necrosis was observed as one of the most distinct pathological features of INH+RIF in liver sections. The hepatocyte swells and then ruptures due to defective osmotic regulation at the cell membrane. Before rupture, membrane blebs form, carrying off cytoplasmic contents (without organelles) into the extracellular compartment [72].

An increase in fibrosis in the T group was observed (Figure 2). This can be explained by the activation of hepatic stellate cell (HSC) and their phenotypic transformation into myofibroblasts, possessing pro-fibrogenic role [76,77]. The transition of HSCs into myofibroblasts is triggered by pro-inflammatory factors like TNFα, IL-1β and IL-6 and pro-fibrogenic factors, especially TGFβ [78,79]. Presence of hemosiderin-laden macrophages (Figure 2) that phagocytosed extravasated red cells in hemorrhagic infarct was observed. The macrophages convert heme iron into hemosiderin, during active red cell breakdown [70]. Accumulation of neutral lipids in hepatocytes results in fatty changes (micro and macrovesicular steatosis) as seen in the section (Figure 2). This may also result in ballooning degeneration [80]. Drug-induced microvesicular steatosis occurs due to primary and secondary mitochondrial dysfunction as a result of energy shortage and inhibition of β-oxidation [81]. A consequent reduction in proton motive force for ATP synthesis is a manifestation of energy deprivation in addition to the deprivation caused by unbalanced ROS generation [82]. The behavior was also reported in previous experiments that showed significant inhibitory action in melittin treated groups when compared to the toxic groups [83,84].

Our results further strengthened the role of melittin in reducing the high rate of lethality, chronic liver injury, attenuating hepatic inflammatory responses and inhibiting hepatocyte apoptosis. Other studies demonstrated that low concentrations of melittin did not have any harmful effects on the tissues and cellular functions [85]. Thus, it is clear from our findings and from the previous work of other researchers that melittin might possess an ameliorative role in biological, biochemical, hematological and histological alterations against hepatotoxicity as a therapeutic drug.

## Conclusions

It is evident from this study that melittin protects isoniazid and rifampicin induced hepatotoxicity. Its use as an anti-inflammatory agent will depend on the development of innovative research protocols to validate its efficacy and safety. Modification of the toxic peptide could overcome the associated ill-effects and pave a way for promoting novel pharmaceutical agents. We propose that melittin could be used as a potential therapeutic agent for attenuating acute liver injury and hepatic failure provided prerequisites are met to evade adverse effects.

## Acknowledgments

The authors would like to thank the Central Research Lab of the Faculty of Science, Sana’a University. In addition, the authors thank Al-Aulaqi Specialized Medical Laboratory and Central Lab of Uni. of Sci. & Tech. Hospital, Sana’a for their support.

## Abbreviations

INH: Isoniazid
RIF: Rifampicin
MEL: Melittin
GSH: Reduced glutathione
DTNB: 1,2 dithio-bis nitrobenzoic acid
TBA: thiobarbituric acid

